# The footprint of colour in EEG signal

**DOI:** 10.1101/2025.10.06.680651

**Authors:** Arash Akbarinia

**Affiliations:** University of Giessen, Germany; Lancaster University Leipzig

## Abstract

Our perception of the world is inherently colourful, reflecting a tight interaction between colour and object processing. Previous studies have shown that colour information can be decoded from non-invasive electroencephalography (EEG) signals when participants view simple, uniformly coloured stimuli. Here, we test whether colour can also be decoded from EEG responses to complex natural images, without explicit colour cueing. We analysed the THINGS EEG dataset, comprising 64-channel recordings from participants viewing 1,800 object concepts (16,740 images), each presented for 100 ms, yielding over 82,000 trials. We established a behavioural ground truth for perceived colour via a psychophysical experiment in which participants selected all perceived colours from a 13-option palette. To model colour-object perception under brief exposure, we used the Segment Anything Model (SAM) to segment images and replaced each segment with its average colour. We then trained an artificial neural network, **CUBE** (**C**olo**U**r and o**B**j**E**ct decoding), with two objectives: (1) alignment with a frozen pretrained CLIP vision encoder, and (2) prediction of human-derived colour distributions. Results show that CUBE robustly decodes colour from EEG signals (average F-score = 0.50). Removing the CLIP alignment reduces performance (F-score = 0.46), indicating that while semantic representations enhance colour decoding, substantial colour information remains decodable from EEG signals even without semantic guidance. Finally, we evaluated object recognition across two EEG and MEG datasets in a 200-class task. Incorporating colour-segmented images improved accuracy by approximately 5% across seven vision encoders. Together, these findings demonstrate that EEG signals during natural vision carry rich colour information that interacts with object processing.

## Introduction

Our visual system makes sense of a scene with remarkable speed. We can attach a simple description such as “green tree” to what we have seen (Potter et al., 2014), in as little as 13 ms. This raises a critical question: what neural representations emerge within such a brief window, and to what extent can they be captured in neuroimaging signals? Here we turn our attention to colour, an effortless and ever-present aspect of vision. Colour not only shapes how we perceive objects (Bramão et al., 2011; Tanaka et al., 2001), but also enhances memorability (Gegenfurtner & Rieger, 2000; Wichmann et al., 2002) and speeds up recognition (Møller & Hurlbert, 1996; Rosenthal et al., 2018).

Colour decoding from neuroimaging has a long history (Paulus et al., 1984; Regan, 1970). Brain activity carries information about chromaticity, luminance, and saturation (Hermann et al., 2022; Pennock et al., 2023; Rozman et al., 2024; Sutterer et al., 2021), the hue circle (Hajonides et al., 2021), the geometry of colour space (Rosenthal et al., 2021), and even unique hues (Chauhan et al., 2023). Fewer studies, however, have examined how colour interacts with object processing. One study suggests that while both shape and colour can be decoded as early as 60-70 ms after stimulus onset, shape-colour congruency emerges later, around 200 ms (Teichmann et al., 2020). These findings are valuable but mostly derive from simplified displays of uniform coloured patches on plain backgrounds. It remains unclear whether colour can be reliably decoded from brain signals when viewing rich, natural scenes– this is the challenge addressed in the present study.

Artificial intelligence and large datasets have recently accelerated progress in decoding. The NSD dataset (Allen et al., 2022), for example, provides large-scale fMRI data covering about 10,000 natural images. Similarly, the THINGS EEG (Gifford et al., 2022) and MEG (Hebart et al., 2023) datasets offer recordings of comparable scale, enabling new opportunities to investigate how natural images are represented in the brain. Alongside these resources, contrastive learning (Radford et al., 2021) has emerged as a powerful tool for decoding. It has already shown strong performance across modalities, from speech recognition (Défossez et al., 2023) to visual object recognition in fMRI (Scotti et al., 2024), EEG (Song et al., 2024), and MEG (Wu et al., 2025).

### CUBE (ColoUr and oBjEct decoding)

We adopted a similar contrastive learning framework on large datasets to investigate colour decoding from brain activity during natural image viewing. We focus on EEG, which, despite its low spatial resolution, offers high temporal resolution, affordability, portability, and the potential for real-time decoding (Benchetrit et al., 2023; Robinson et al., 2023). Rather than collecting a new dataset, we augmented the THINGS EEG dataset (Gifford et al., 2022) with colour annotations from a largescale psychophysical experiment matched to the original conditions. In a rapid serial visual presentation (RSVP) paradigm, participants viewed each image for 100 ms and selected all perceived colours from a 13-option palette. (see Figure 1, panel C).

**Figure 1:**
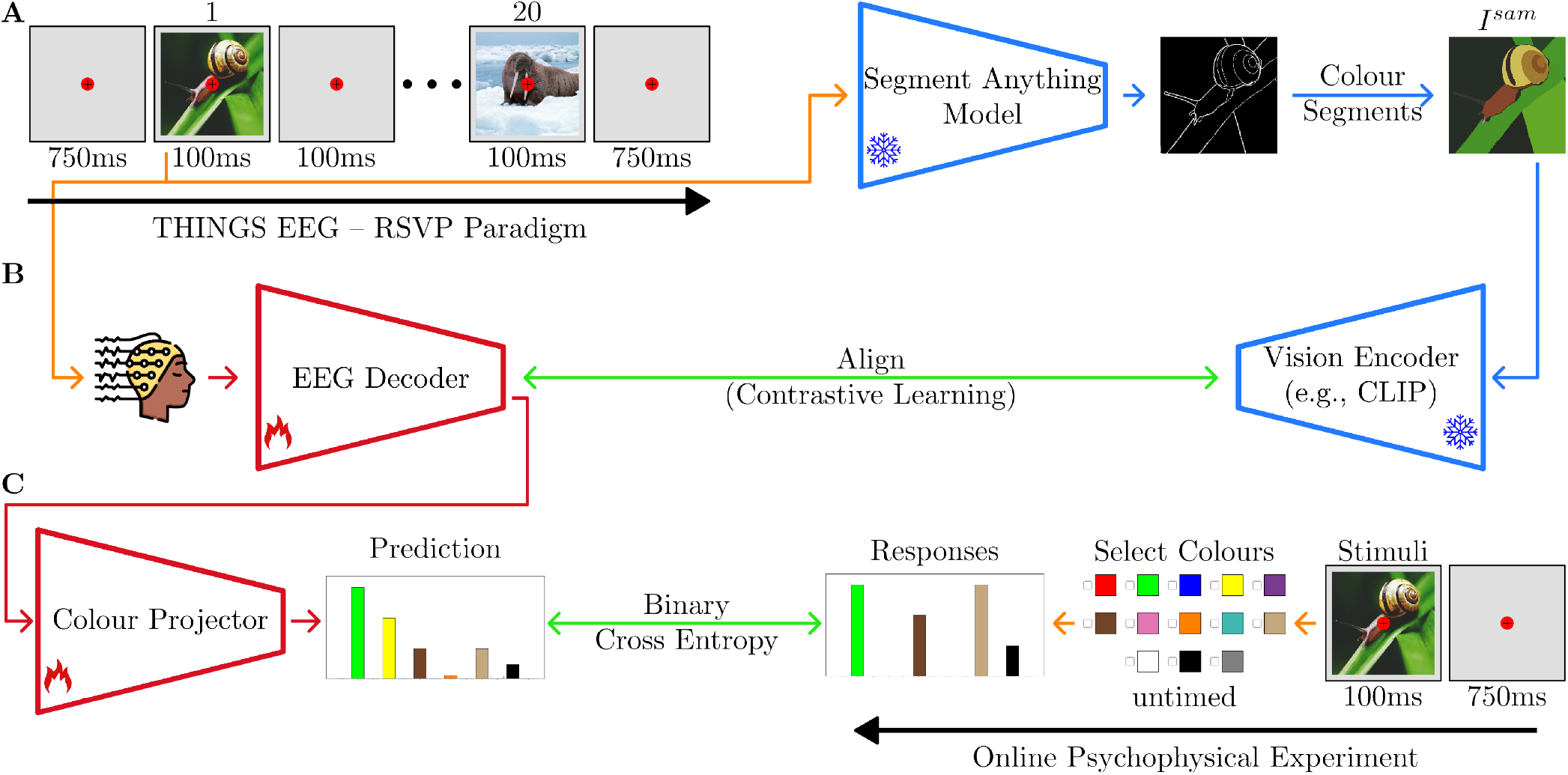
The overview of **C**olo**U**r and o**B**j**E**ct decoding–**CUBE. A**: The RSVP paradigm used to collect the THINGS EEG dataset (Gifford et al., 2022). **B**: An EEG decoder aligns brain activity with features from a pretrained vision encoder applied to colour-segmented images. **C**: A linear projection layer maps the aligned representation onto the behavioural colour responses.

We trained a lightweight EEG decoder–a simple artificial neural network with two linear layers and a residual connection–to align EEG representations with features extracted from a frozen CLIP visual encoder, a pretrained vision-language model (Radford et al., 2021). To better capture perceptual colour structure, which is organised at the level of objects rather than individual pixels (Gegenfurtner, 2025), we processed each image (*I*^*org*^ ) using the Segment Anything Model (SAM) (Kirillov et al., 2023). For each segmented region, pixel colours were averaged to generate uniformly colour-segmented images (*I*^*sam*^ ) (see Figure 1, panel B), providing a closer approximation to colour perception under brief viewing.

We added a linear projection layer to the EEG decoder to produce a 13-dimensional output predicting behavioural colour responses. Given the multi-class nature of the task, decoding performance was evaluated using the F-score. The full CUBE model achieved an average F-score of 0.50 across participants. Ablating the CLIP alignment, thereby removing access to semantic information, reduced performance to an F-score of 0.46. This reduction highlights the interaction between object and colour processing, while still indicating that substantial colour information remains decodable from EEG signals, well above chance (0.17) and approaching the noise ceiling defined by average human agreement (0.64) in this rapid RSVP paradigm. Together, these results suggest that EEG signals recorded during brief viewing of complex natural images carry robust, decodable colour information.

Encouraged by this observation–that semantic object information influences colour decoding, consistent with findings from both behavioural (Bramão et al., 2011; Gegenfurtner, 2025) and neural literature (Rosenthal et al., 2018; Tanaka et al., 2001)–we hypothesised that this relationship is bidirectional, incorporating colour information into the contrastive alignment framework should conversely improve object decoding.

To test this, we employed the Uncertainty-aware Blur Prior algorithm (Wu et al., 2025), aligning CLIP features with both foveation-blurred original images (*I*^*org*^ ) and colour-segmented images (*I*^*sam*^ ) (see Figure 2). Results show that CUBE improves state-of-the-art object decoding by a consistent 5% across all participants in both EEG and MEG datasets. Critically, repeating the analysis with greyscale versions of *I*^*sam*^, which preserves segmentation while removing colour information, abolished this performance gain, underscoring the importance of colour-object interactions for decoding.

**Figure 2:**
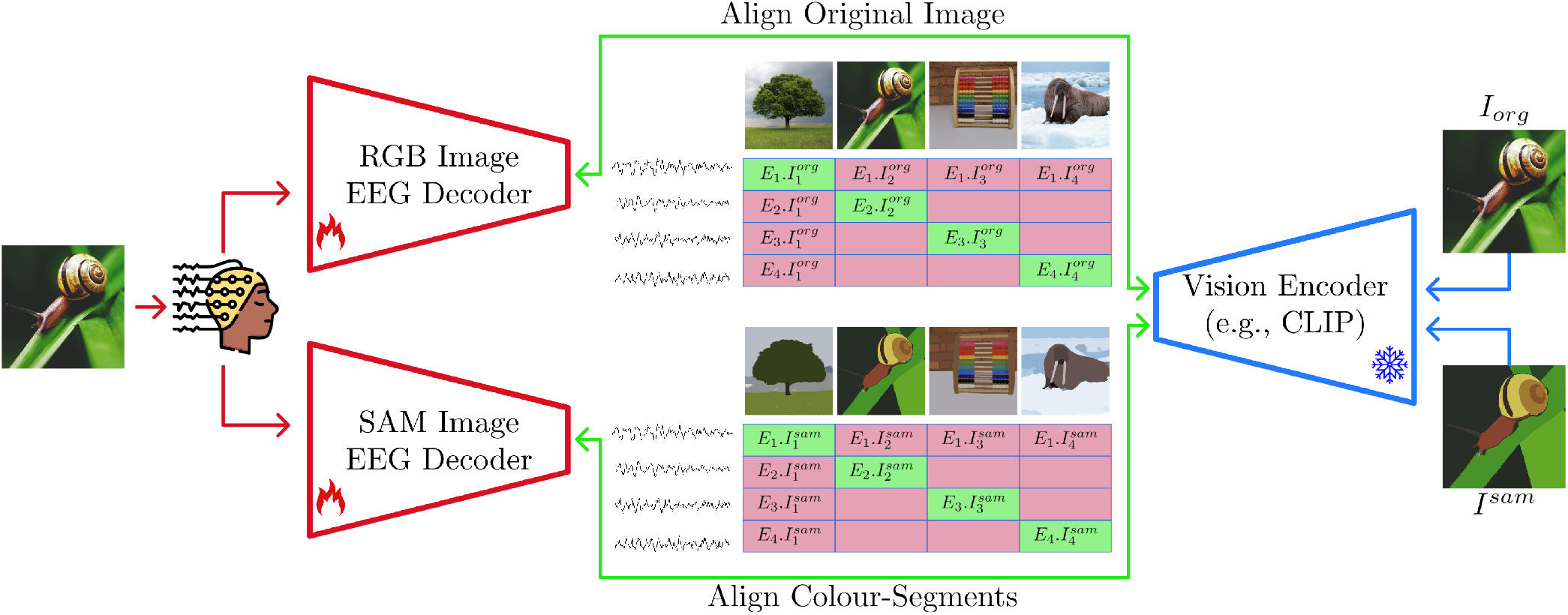
Object decoding using two EEG decoders trained to align with (1) the original RGB images viewed by participants and (2) colour-segmented versions of the same images generated using the Segment Anything Model (SAM) (Kirillov et al., 2023). Colour-segment alignment follows the standard contrastive learning (Radford et al., 2021), whereas RGB image alignment employs the Uncertainty-aware Blur Prior algorithm (Wu et al., 2025).

## Method

We primarily focused on the THINGS EEG dataset and, secondarily, on the MEG dataset, both derived from a subset of the THINGS collection (Hebart et al., 2019), a high-quality set comprising 1,854 diverse object concepts. We generated colour annotations for the images through an online psychophysical experiment and employed a contrastive learning framework to train our networks ^1^.

### Neuroimaging datasets

#### THINGS EEG (Gifford et al., 2022)

Recordings were collected from 10 participants using an RSVP paradigm (Intraub, 1981), where each image was shown for 100 ms, followed by a 100 ms blank (Figure 1, Panel A). EEG was recorded with a 64-channel cap. The training set included 1,654 concepts (10 images per concept, 4 repetitions per image), and the test set 200 unseen concepts (1 image per concept, 80 repetitions per image). Preprocessing followed the original paper: signals were epoched 0-1000 ms post-stimulus, downsampled to 250 Hz, and reduced to 17 occipito-parietal channels most relevant to vision^2^. Repetitions were averaged to enhance the signal-to-noise ratio, resulting in 16,540 training and 200 test samples per participant.

#### THINGS MEG (Hebart et al., 2023)

Recordings from 4 participants with 271 channels, each image presented for 500 ms followed by a 1000 ± 200 ms interval. The training set included 1,854 concepts (12 images per concept, 1 repetition each), and the test set comprised 200 concepts (1 image per concept, 12 repetitions each). As with the EEG dataset, test concepts were excluded from training to facilitate zero-shot evaluation. Preprocessing mirrored the EEG pipeline, with MEG signals downsampled to 200 Hz.

### Psychophysical experiment

When an image is presented for only 100 ms, a key question is how much of the scene can be consciously accessed and understood (Keysers et al., 2001). To approximate participants’ colour perception in the EEG experiment, we conducted a psychophysical study with a similar presentation rate. Each trial began with a fixation cross (750 ms), followed by the image (100 ms). Participants, guided by written instructions, were asked to select all perceived colours from a 13-colour palette without linguistic labels (Figure 1, Panel C). Eleven colours were derived from universal colour terms shared across cultures (Berlin & Kay, 1991), with two additional colours (turquoise and beige) included due to their frequent usage (Mylonas & MacDonald, 2016).

To annotate the full THINGS EEG dataset (16,740 images), the experiment was conducted via Prolific (Palan & Schitter, 2018) with 133 compensated participants from diverse backgrounds. Each participant viewed approximately 450 images. Training trials were limited to a 5-second response window due to the high stimulus volume, while test trials were unconstrained to ensure comprehensive colour reporting; however, average response latencies remained comparable across both conditions.

In this RSVP-style paradigm (100 ms exposure), participants typically recalled few colours (mean: 2.1 per image). Reports exhibited a bias toward salient fore-ground objects, whereas background colours were less frequently reported unless they occupied large, uniform regions. Beyond annotating the THINGS EEG dataset, this publicly released large-scale study provides a valuable resource for investigating colour perception under brief, ecologically valid conditions.

### Visual neural decoding

To decode colour and object from neuronal data, we adopted a contrastive learning paradigm (Radford et al., 2021), which has been widely used in neuroimaging decoding research (Li et al., 2024; Scotti et al., 2024; Song et al., 2024). In this framework, an EEG decoder is trained to align its output representations with those of a pretrained vision encoder, often a variant of CLIP (Figure 2). Similar strategies have also been applied with other modalities, such as language encoders (Akbarinia, 2024) and depth encoders (Zhang et al., 2025).

Formally, let 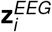 denote the feature representation predicted by the EEG decoder for sample *i*, and 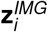 the corresponding representation from the vision encoder. The goal of contrastive learning is to maximise similarity between matching pairs 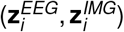 while minimising similarity with all non-matching pairs in the batch. This is achieved with a symmetric cross-entropy objective:

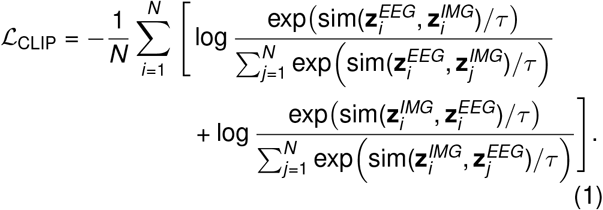

where *N* is the batch size, *τ* is a learnable temperature parameter, and sim(·, ·) represents a similarity function. The objective is implemented using a categorical cross-entropy loss. This formulation encourages EEG and image representations of the same stimulus to be close in the embedding space, while pushing apart representations corresponding to mismatched pairs.

One persistent challenge in neuroimaging applications is limited dataset size: current datasets are relatively small for deep learning, often leading to overfitting during training and poor generalisation at test time. A recently proposed technique, the *Uncertainty-aware Blur Prior* (Wu et al., 2025), mitigates this issue by applying a foveated blur to the original images *I*^*org*^, approximating how participants perceive visual stimuli. By attenuating high-frequency details, this approach reduces a key source of overfitting. We adopt this strategy in our training framework by applying the foveation procedure described in Wu et al. (2025) to *I*^*org*^ .

Building on this idea, we hypothesised that colour-segmented images more closely reflect participants’ perceived colours and their associations with objects under brief exposures (100 ms). To evaluate this, we conducted pilot experiments in which participants viewed *I*^*org*^ for 100 ms, followed by a 750 ms grey screen. Subsequently, *I*^*sam*^ was presented either alone or alongside a greyscale version of *I*^*org*^ . Participants were instructed to identify pixels whose colours were inconsistent with the originally viewed scene. Only a small number of mismatches were reported, suggesting that colour-segmented images provide a close approximation of perceived colours, objects, and their associations under such brief viewing conditions.

Motivated by these findings, we extended our object decoding framework by jointly aligning a second EEG decoder with CLIP features extracted from colour-segmented images, denoted as *I*^*sam*^ . These images are derived from the original input *I*^*org*^ using SAM-1 (Kirillov et al., 2023) with default global segmentation parameters, except for an increased resolution of 64 points per side and a stability score threshold of 0.92. We then introduce a contrastive loss term ℒ_CLIP_ between 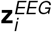 and 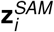 to support object decoding in CUBE.

The *EEG Decoder* in CUBE follows Wu et al. (2025), consisting of two linear layers with GELU activation and a residual connection. The *Colour Projector* comprises two linear layers with ReLU activation, mapping CLIP-aligned features to a 13-colour palette. We utilised OpenCLIP (Cherti et al., 2023) to extract features across various vision encoders. Networks were trained for 50 epochs with a batch size of 1024 using the AdamW optimiser (Loshchilov & Hutter, 2017), with a learning rate of 1 *×* 10^−4^ and weight decay of 1 *×* 10^−4^. All other configurations followed Wu et al. (2025).

For each experiment, two training regimes were considered. *Intra-participant* networks were trained and evaluated on data from the same participant, whereas *inter-participant* networks were trained on all but one participant, with the held-out participant used for testing.

## Results

### Colour decoding

Colour decoding is inherently a multi-class task, as multiple colours may co-occur within a single image. Unlike object recognition, colour perception shows striking individual differences, particularly under brief viewing (Bosten, 2022; Mollon et al., 2017). For example, one participant may label *wood* as *brownish*, whereas another may choose a more *beige* shade (Lafer-Sousa et al., 2015). To accommodate this multi-class structure and the variability across observers, we quantified agreement between two human responses using the F-score:

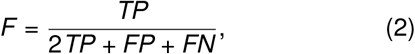

where *TP, FP*, and *FN* denote true positives, false positives, and false negatives, respectively. We chose the F-score over the closely related Jaccard index (Jaccard, 1901), another metric for set comparison, because the F-score places greater emphasis on true positives (*TP*s). This property is better suited to assessing colour fidelity, as it prioritises dominant colours consistently selected by participants.

We directly compared behavioural and neural data by applying the same metric to quantify agreement between EEG-decoded colours and the average human responses. As neither the human averages nor the model predictions are binary, both were thresholded prior to evaluation. All reported F-scores use a threshold of 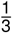, based on the rationale that at least one-third of participants must agree on a given colour for an image.

The results of colour decoding on the THINGS EEG dataset are shown in Figure 3. Overall, the CUBE model achieves an F-score above 0.50, approaching the noise ceiling (0.64)–defined as the average agreement among participants–and substantially exceeding chance (0.17), estimated over 10,000 iterations using two randomly selected colours per trial to match typical participant responses. Performance also surpasses a distribution-aware baseline (0.23), computed by sampling colours from the training-set distribution. Inter-participant models achieve lower F-scores (0.33) but remain well above chance, indicating a shared representation of colour across individuals (Gegenfurtner, 2003).

**Figure 3:**
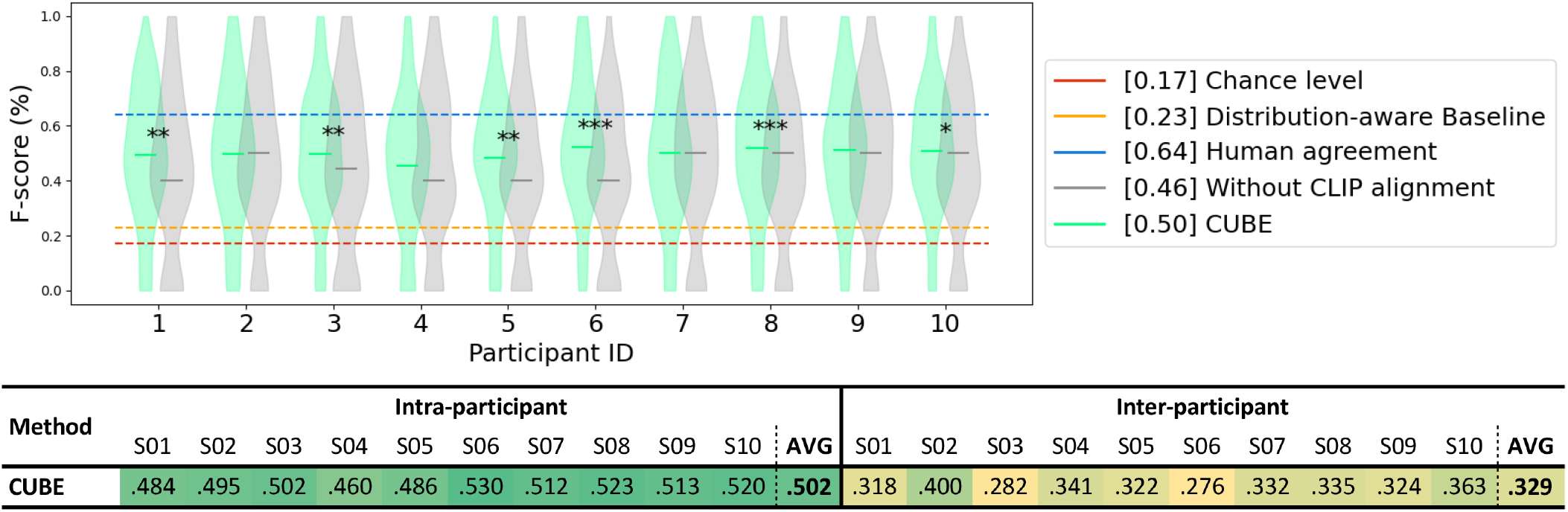
Colour decoding performance of CUBE. **Top**: F-score distributions over 200 test images. Chance level is shown in red and the noise ceiling from behavioural agreement in blue. Asterisks mark significant differences between models with and without CLIP alignment. **Bottom**: Average F-scores for within- (intra) and across-participant (inter) training. Table cells are colour-coded from green to yellow as F-scores decline.

Ablating CLIP alignment during training–thereby removing semantic guidance–yields an F-score of 0.46, still markedly above chance. This demonstrates that the network can extract meaningful colour information directly from EEG signals without relying on external semantic features. Nevertheless, for most participants, colour decoding improves significantly (Students *t*-test) when CLIP alignment is included, consistent with evidence that object and semantic information modulate colour perception (Gegenfurtner, 2025; Tanaka & Presnell, 1999; Witzel & Gegenfurtner, 2018). Although the reported results are based on the CoCa-ViT-L-14 architecture (Yu et al., 2022), no significant differences were observed when using alternative architectures.

The F-scores vary by 7% between the best and worst participants (53% for participant 06 and 46% for participant 04), yet the distributions across the test set appear qualitatively similar. This consistency likely reflects the fact that the colour ground truth is based on average human responses rather than each participants individual colour perception during the EEG experiment (Bosten, 2022). Consequently, higher decoding accuracy might be achievable if the behavioural ground truth corresponded to the neural data of the same individual.

The examples in Figure S1 show that neurally decoded colours often remain plausible even in trials with low F-scores. For instance, the *Flax Seed* image is behaviourally labelled *brown-beige*, while the decoded hues are neighbouring *orange-red*, indicating only a small mismatch. Similar swaps between red and orange appear for the *Omelette* and *Fruit* images–a known challenge even for computer vision models (Parraga & Akbarinia, 2020). In other cases, such as the *Elephant*, mechanisms like colour constancy (Akbarinia & Parraga, 2017; Gil Rodríguez et al., 2024) or memory (Hansen et al., 2006) may drive participants to report grey despite physically *yellowish-beige* pixels, reflecting processes requiring longer temporal integration. Overall, the neurally decoded colours qualitatively appear both meaningful and perceptually coherent.

### Object decoding

Object decoding in the THINGS EEG and MEG datasets is formulated as a retrieval task. Each network is evaluated in a zero-shot setting, where, among 200 candidate images, the one with the highest cosine similarity to the decoded EEG features is taken as the predicted object category.

Table 1 reports the object decoding accuracy of CUBE and several comparison models on the **THINGS EEG dataset** (Gifford et al., 2022). CUBE achieves 57% top-1 and 86% top-5 accuracy, representing a 6% improvement over UBP (Wu et al., 2025). This gain is consistent across all participants, indicating robust decoding boost. Accuracy peaks at 66% top-1 and 93% top-5 for the best participant–remarkable given the inherently noisy nature of EEG signals. In inter-participant evaluation, the improvement is more modest–just over 1% in both top-1 and top-5 accuracy–reflecting the substantial challenges of cross-subject decoding, including variability in neural responses (Wei et al., 2021) and individual differences in visual processing (De Haas et al., 2019).

**Table 1:**
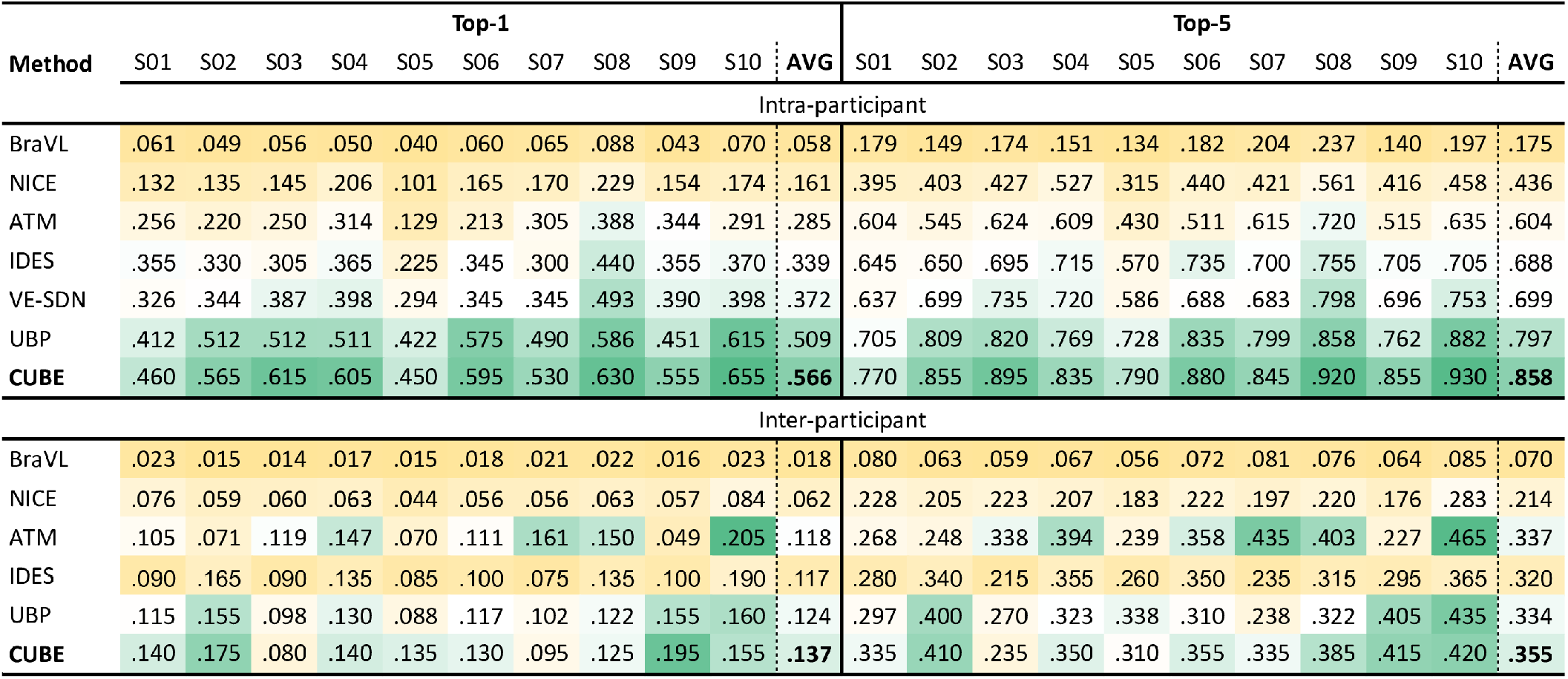
Object decoding performance of CUBE on the THINGS EEG dataset (Gifford et al., 2022) across 200 object categories. Comparison methods from the literature include BraVL (Du et al., 2023), NICE (Song et al., 2024), ATM (Li et al., 2024), IDES (Akbarinia, 2024), VE-SDN (Chen et al., 2024), and UBP (Wu et al., 2025). Table cells are colour-coded from green to yellow as accuracies decrease.

We subsequently assessed whether the performance gains of CUBE generalise across diverse visual architectures. As illustrated in Figure 4, CUBE yields a statistically significant 5% increase in object decoding accuracy across seven distinct OpenCLIP vision encoders (Cherti et al., 2023). To determine if this enhancement stems from the semantic segmentation structure of *I*^*sam*^ rather than intrinsic colour features, we evaluated CUBE variants aligned with EEG using greyscale *I*^*sam*^ . These control models–where colour information is absent but semantic structure is preserved–yielded only a marginal 1% average gain, which was inconsistent across encoders and resulted in performance degradation for the CoCa-B32 architecture. Collectively, these findings demonstrate that colour provide a substantive and robust contribution to object decoding. This aligns with the observation that under rapid stimulus exposure (100 ms), colour is among the earliest visual attributes to be processed, leaving a salient and quantifiable signature in the neural signal.

**Figure 4:**
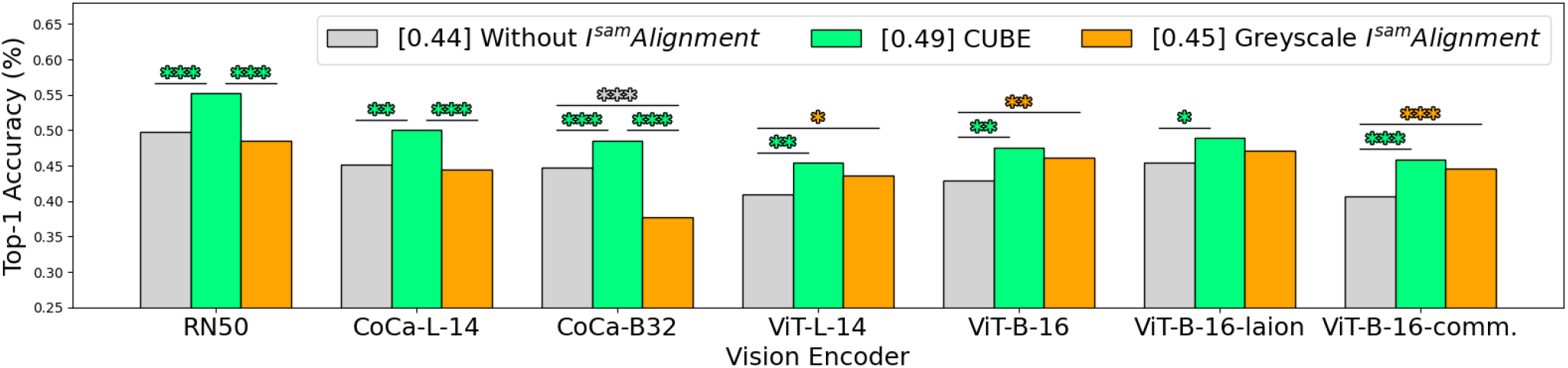
Impact of colour modelling on object decoding. Decoding performance across seven vision encoders on the THINGS EEG dataset (Gifford et al., 2022), evaluated over 200 object categories. All encoders were pretrained using OpenCLIP (Cherti et al., 2023). Asterisks indicate statistically significant differences between conditions.

Table 2 reports object decoding accuracy for CUBE and two comparison models on the **THINGS MEG dataset** (Hebart et al., 2023). The results closely parallel the EEG findings: CUBE improves intra-participant accuracy by roughly 5% for both top-1 and top-5, and inter-participant accuracy by about 1%. These findings demonstrate that the decoding boost provided by colour-segmented images generalises beyond EEG to other neuroimaging modalities.

**Table 2:** Object decoding performance of CUBE on the THINGS MEG dataset (Hebart et al., 2023) across 200 object categories. Comparison methods from the literature include BraVL (Du et al., 2023), NICE (Song et al., 2024), and UBP (Wu et al., 2025). Table cells are colour-coded from green to yellow as accuracies decrease.

| Method | Top-1 |  |  |  |  | Top-5 |  |  |  |  |
| --- | --- | --- | --- | --- | --- | --- | --- | --- | --- | --- |
|  | S01 | S02 | S03 | S04 | AVG | S01 | S02 | S03 | S04 | AVG |
| Intra-participant |  |  |  |  |  |  |  |  |  |  |
| NICE | .096 | .185 | .142 | .090 | .128 | .278 | .478 | .416 | .266 | .360 |
| UBP | .150 | .460 | .273 | .185 | .267 | .380 | .805 | .590 | .435 | .552 |
| CUBE | .165 | .520 | .335 | .235 | .314 | .435 | .850 | .625 | .480 | .598 |
| Inter-participant |  |  |  |  |  |  |  |  |  |  |
| UBP | .020 | .015 | .027 | .025 | .022 | .057 | .172 | .100 | .080 | .104 |
| CUBE | .020 | .050 | .035 | .025 | .033 | .075 | .175 | .100 | .105 | .114 |

## Discussion

EEG, whose core technology dates back nearly a century (Berger, 1929), measures tiny fluctuations in ionic potentials to non-invasively record brain activity. The resulting signal is notoriously noisy and has low spatial resolution, reflecting the aggregate activity of billions of neurons (Azevedo et al., 2009; Goriely, 2025). Despite these limitations, EEG has long been an invaluable tool– both clinically and for advancing our understanding of the brain. Recent work suggests that the decoding capabilities of EEG, and neuroimaging more broadly, are undergoing a major leap forward, enabled by AI and large-scale datasets. In particular, EEG benefits from its exceptionally high temporal resolution. We can now decode speech from three seconds of EEG with remarkable accuracy (Défossez et al., 2023), and in the visual domain, emerging work is progressing toward 3D object reconstruction (Guo et al., 2025) and even video decoding (Liu et al., 2024).

Here, we show that object decoding reaches a remarkable 57% accuracy–far above the 1/200 chance level–using just one second of EEG data. Likewise, we demonstrate for the first time that perceived colours in complex natural images can be decoded with high reliability (F-score = 0.5). Strikingly, EEG recorded during natural viewing–without any explicit colour cues–can recover colours with reliability approaching that of average behavioural responses. One might expect decoding performance to improve further if neural and behavioural data were collected from the same individuals, thereby avoiding the alignment problem (Robinson et al., 2023) and enabling models to more precisely capture individual differences in perception and colour experience (Bosten, 2022; De Haas et al., 2019).

### The interaction between colour and object

Although the interaction between colour and object processing remains surprisingly little understood (Gegenfurtner, 2025; Taylor & Xu, 2021; Teichmann et al., 2020), substantial evidence suggests that colour information plays an important role in object recognition (Bramão et al., 2011; Rosenthal et al., 2018), and, conversely, object and scene semantics are known to influence perceived colours (Akbarinia, 2025b; Bloj et al., 1999; Hansen et al., 2006). Our decoding results for both colour and objects further support this bidirectional relationship at the level of neural representations: object alignment improves colour decoding by 4%, while incorporating colour-segmented information enhances object decoding by 5%.

A central question in visual cognitive neuroscience is whether colour is encoded first–with object boundaries emerging later, as in a bottom-up region-growing process–or whether objects are parsed first and colours subsequently filled in, reflecting a more top-down mechanism (Palmer, 1981). To investigate this, we compared colour and object decoding within CUBE.

A direct comparison, however, is not straightforward. First, the evaluation metrics differ: F-score for multiclass colour decoding versus accuracy for single-label object decoding. Second, the ground truths differ: object labels are objective (“this is a snail”), whereas colour annotations reflect subjective averages across observers. Third, colour and object are tightly intertwined, introducing potential confounds. Colour-diagnostic objects can facilitate colour decoding (e.g., bananas are yellow), while natural colour statistics can bias object recognition (Tanaka & Presnell, 1999; Therriault et al., 2009). For example, a uniformly orange sphere may be decoded as an orange, whereas a striped orange sphere may instead be classified as a volleyball. Fourth, material properties introduce additional complexity (Schmidt et al., 2025), as certain materials (e.g., wood) exhibit characteristic colour–texture associations.

Despite these methodological challenges, within our framework colour decoding performs relatively better when normalised to distribution-aware chance level and noise ceilings: the colour-decoding F-score reaches 0.66 on a 0–1 scale, exceeding the object-decoding accuracy of 0.57.

To examine temporal dynamics, we trained models on 100 ms EEG segments from successive intervals, e.g., [0, 100), [100, 200), [900, 1000). To estimate the minimal temporal information required for above-chance decoding, we also evaluated shorter windows extending from stimulus onset up to time *t*, e.g., [0, 25) ms. As shown in Figure 5, peak decoding accuracy for both colour and object occurs for most participants in the [200, 300) ms interval, and for a smaller subset in [100, 200) ms. This suggests that the most robustly decodable representations emerge shortly after stimulus offset.

**Figure 5:**
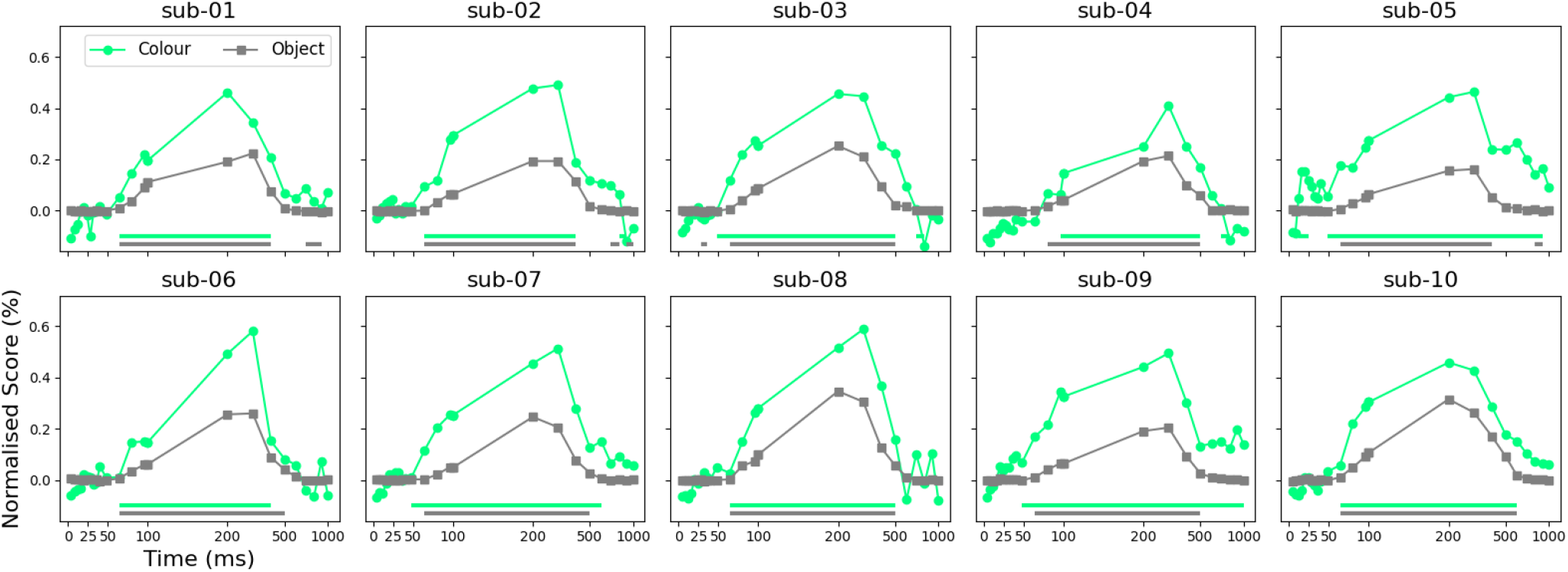
Colour and object decoding across temporal epochs. The x-axis (0 ms) marks stimulus onset; the y-axis shows normalised F-scores (colour) and top-1 accuracy (object), scaled by distribution-aware chance level and noise ceiling. Data points after 100 ms use EEG intervals [*t* − 100, *t* ), and points before 100 ms from [0, *t* ). Significance lines indicate intervals with decoding above chance (*p* < 0.01).

A comparison of the time courses of statistical significance for colour and object decoding reveals that colour decoding reaches significance earlier than object decoding–on average at 70 ms for colour versus 77 ms for objects. These timeframes are broadly consistent with previous findings in the literature (Teichmann et al., 2020). However, in contrast to that study, we observe an earlier onset of significant decoding for colour, a pattern that is consistent across participants.

Taken together, these results suggest that colour may play a more prominent role during the earliest stages of visual processing (Gegenfurtner, 2003), aligning with the subjective impression that colour rapidly “pops out”. At the same time, this difference remains an open and interesting question that warrants further investigation.

### Limitations of the EEG decoding

What are the limitations of current EEG decoding frameworks? A major challenge is the substantially lower cross-participant performance compared with within-participant decoding, driven by large individual differences in visual processing across both sensory and perceptual levels (Bosten, 2022; De Haas et al., 2019). Signal quality is further influenced by technical factors such as electrode placement and impedance, which can vary across sessions and participants. One promising direction is to pretrain models on large, diverse EEG datasets (Huang et al., 2025), analogous to large language models, and then fine-tune them for individual participants. This strategy may help bridge the gap between generalisation and personalisation, and could be particularly valuable for practical neuroimaging applications, such as brain-machine interfaces for individuals with severe motor impairments (Chaudhary et al., 2015).

## Conclusion

In this article, we introduced **CUBE** (**C**olo**U**r and o**B**j**E**ct decoding) and highlighted the importance of jointly representing colour and object features in neuroimaging decoding. Our results show that EEG signals contain reliable, decodable colour information–even during object recognition tasks with no explicit colour cues and under very brief viewing conditions (100 ms). We further demonstrated that incorporating colour-segmented information into a standard contrastive-learning alignment framework boosts object decoding by about 5% across participants in both EEG and MEG. Overall, our findings open a novel avenue for future work: theoretically, enabling the investigation of individual colour perception in more ecological settings by decoding colour from neuroimaging signals under naturalistic viewing; and practically, offering potential benefits for applications in braincomputer interfaces and clinical psychology.

## Acknowledgements

This research was funded by the Deutsche Forschungsgemeinschaft SFB/TRR 135 (grant number 222641018) TP S. Parts of this work were presented in abstract form at the European Conference on Visual Perception and the Cognitive Computational Neuroscience (Akbarinia, 2025a).

## Qualitative examples

**Figure S1:**
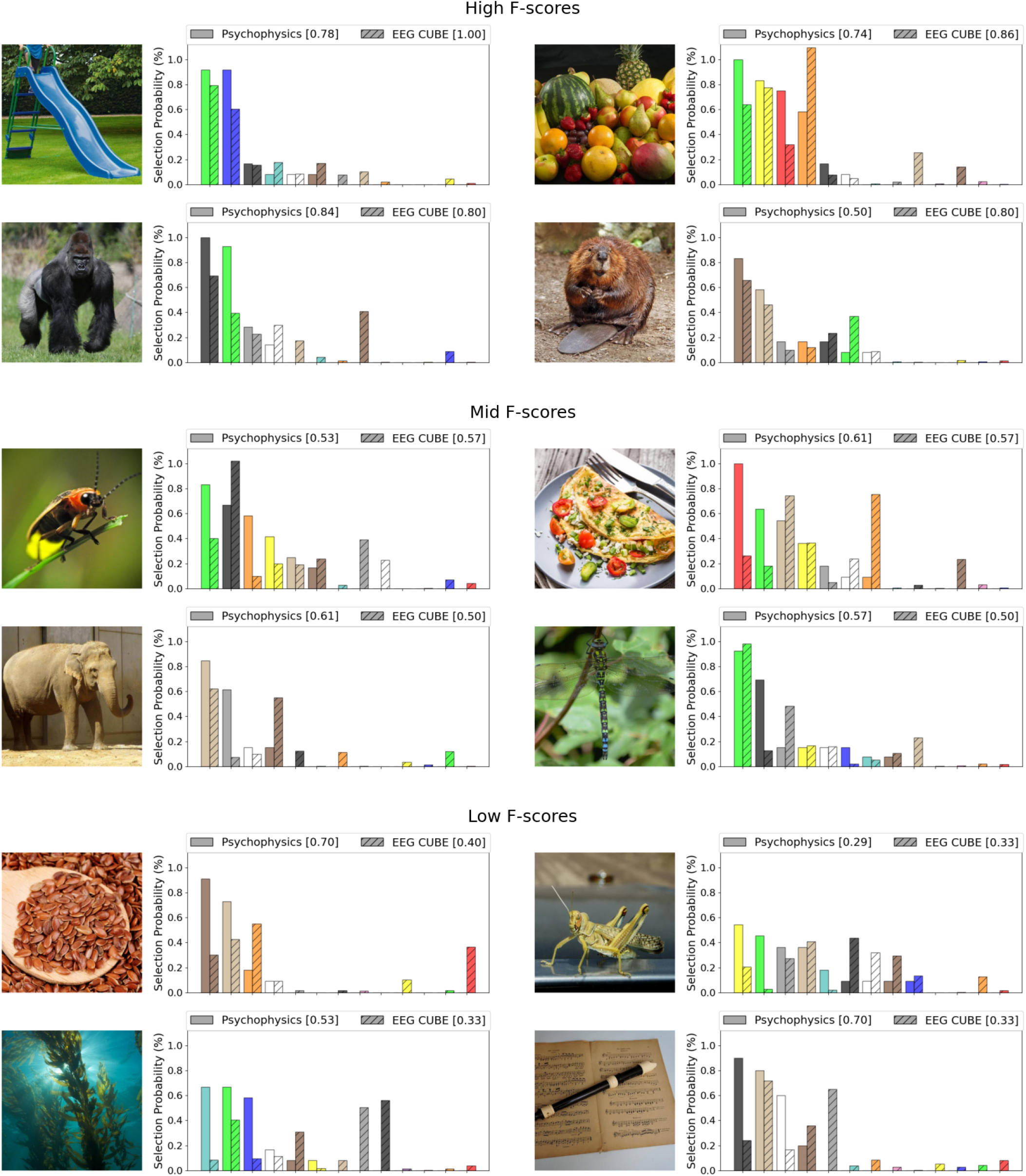
Examples of test images alongside the corresponding colour selections made by human participants in a psychophysical experiment and the colours decoded by CUBE from EEG signals. The reported F-score for the psychophysical data reflects the average inter-participant agreement computed using a leave-one-out strategy, whereas the EEG CUBE F-score represents the agreement between the models predictions and the average human selections.

## Footnotes

1 The experimental materials are available publicly on https://arashakbarinia.github.io/projects/cube/.

2 P7, P5, P3, P1, Pz, P2, P4, P6, P8, PO7, PO3, POz, PO4, PO8, O1, Oz, O2

